# Investigating cross-memory interactions of timing structures

**DOI:** 10.64898/2026.01.10.698757

**Authors:** Sophie Thong, Joshua Hendrikse, Trevor T. -J. Chong, James P. Coxon

## Abstract

Motor and episodic memories are classically conceptualised as distinct memory domains. However, both memory sequences are characterised by order and timing of their elements. Generalisation has been previously documented between these different memories, with a shared ordinal structure facilitating learning. However, it is unclear whether this generalisation also applies to the timing structure of a sequence. In a between-subject design, healthy adults (*N* = 143) learned and reproduced motor and episodic memory sequences with either the same or different timing structures. Contrary to expectations, recall of an episodic timeline was poorer when preceded by learning a motor sequence with the same timing structure. Motor sequence reproduction was not affected by prior episodic encoding. The findings do not support the hypothesis of cross-domain generalisation of timing structure between episodic and motor memories. However, the observed interference may suggest some overlap in the mechanism encoding timing information between memory systems.

## Introduction

Classical models of memory posit that memories of skills and actions (motor memories) and of daily events (episodic memories) are supported by distinct neural systems and mechanisms (Henke, 2010; Squire, 2004). However, action and event memory sequences are similarly characterised by the order and timing (durations) of their elements (Evensmoen et al., 2025; Kornysheva et al., 2013). Interestingly, studies have documented bi-directional generalisation (i.e., facilitated learning and recall of a subsequent memory) between motor and declarative sequences when they shared an ordinal structure (Failla et al., 2025; Mosha & Robertson, 2016). This process of generalisation is deemed to occur through overlapping neural ensembles (Herszage & Censor, 2018; Robertson, 2022; Schlichting & Frankland, 2017), and thus suggests that structural aspects of memories (e.g., order and timing) may be encoded by shared neural networks. However, it remains unknown whether cross-memory generalisation occurs on the basis of a shared timing structure between different memories.

In the timing literature, timing information has been shown to be generalisable across motor and perceptual domains. Specifically, training participants’ acuity in perceiving the durations of stimuli has been shown to facilitate performance of subsequent actions within similar duration ranges (Meegan et al., 2000; Sanchez-Moncada et al., 2024). Similar facilitative effects have been found for perceptual timing when motor timing is trained first (Guo et al., 2025). These findings support the hypothesis that timing information is generally encoded by the motor system, as motor regions have been implicated across both motor and perceptual timing tasks (Breska & Ivry, 2016; Merchant et al., 2013; Sanchez-Moncada et al., 2024).

Yet findings of generalisation between motor and perceptual domains cannot be readily extrapolated to the episodic domain, as episodic timing is encoded and recalled differently from perceptual timing. Perceptual tasks often involve active or anticipatory estimation of the durations of one or two stimuli as they occur (Breska & Ivry, 2016), while in episodic timing, the timing of sequential events are not actively attended to but reconstructed once events have concluded (MacDonald, 2014; Tsao et al., 2022).

Nonetheless, one recent study found activation of motoric regions during episodic timing recall, which suggests that the motor system may also encode timing information across memory domains (Evensmoen et al., 2025). Together, it remains unclear whether sequence timing information is generalisable across motor and episodic memories.

This study investigated whether generalisation between motor and episodic memory sequences occurs on the basis of a shared timing structure between both sequences.

Extrapolating from past findings of ordinal cross-memory generalisation (Mosha & Robertson, 2016; Mutanen et al., 2020), we hypothesised that participants would demonstrate better recall for the timing structure of a motor sequence if they had first encoded an episodic sequence with the same timing structure, and vice versa when the order of task completion was reversed.

## Methods

### Pre-registration disclosures

This study was the second component of a study pre-registered on the Open Science Framework (OSF) on 13 June 2024 (OSF pre-registration).

### Participants

One hundred and forty-three healthy, right-handed participants (M_age_ = 23.43 ± 4.65 years, 36 male) were recruited through convenience sampling in Melbourne, Australia.

Participants did not have a current diagnosis of psychiatric or psychological illness, and provided their written informed consent to participate in the study. This study was approved by the Monash University Human Research Ethics Committee (Project ID: 38080).

### Design

Participants completed two separate tasks probing episodic and motor memory, in counterbalanced order. To test for cross-domain generalisation of temporal information, we manipulated the timing structure across the two tasks, which were either matched (‘same’) or unmatched (‘different’). This study therefore adopted a 2 x 2 factorial design, with the between-subjects factors of ‘Timing structure’ (same vs. different), and Order of task completion (‘episodic task followed by motor task vs. motor task followed by episodic task’) (Figure 1a). Participants were randomly allocated to one of four groups:

- EM-S Group who completed *the episodic task first* (E), followed by the *motor task*
- (M) with the *Same* (S) timing structure between tasks (*N* = 30, 9 male)
- EM-D Group who completed the *episodic task first* (E), followed by the *motor task*
- (M) with Different (D) timing structure between both tasks (*N* = 30, 8 male)
- ME-S Group who completed the *motor task first* (M), followed by the *episodic task*
- (E) with the *Same* (S) timing structure between tasks (*N* = 30, 7 male)
- ME-D Group who completed the *motor task first* (M), followed by the *episodic task*
- (E) with the *Different* (D) timing structure between tasks (*N* = 33, 8 male)

**Figure 1.**
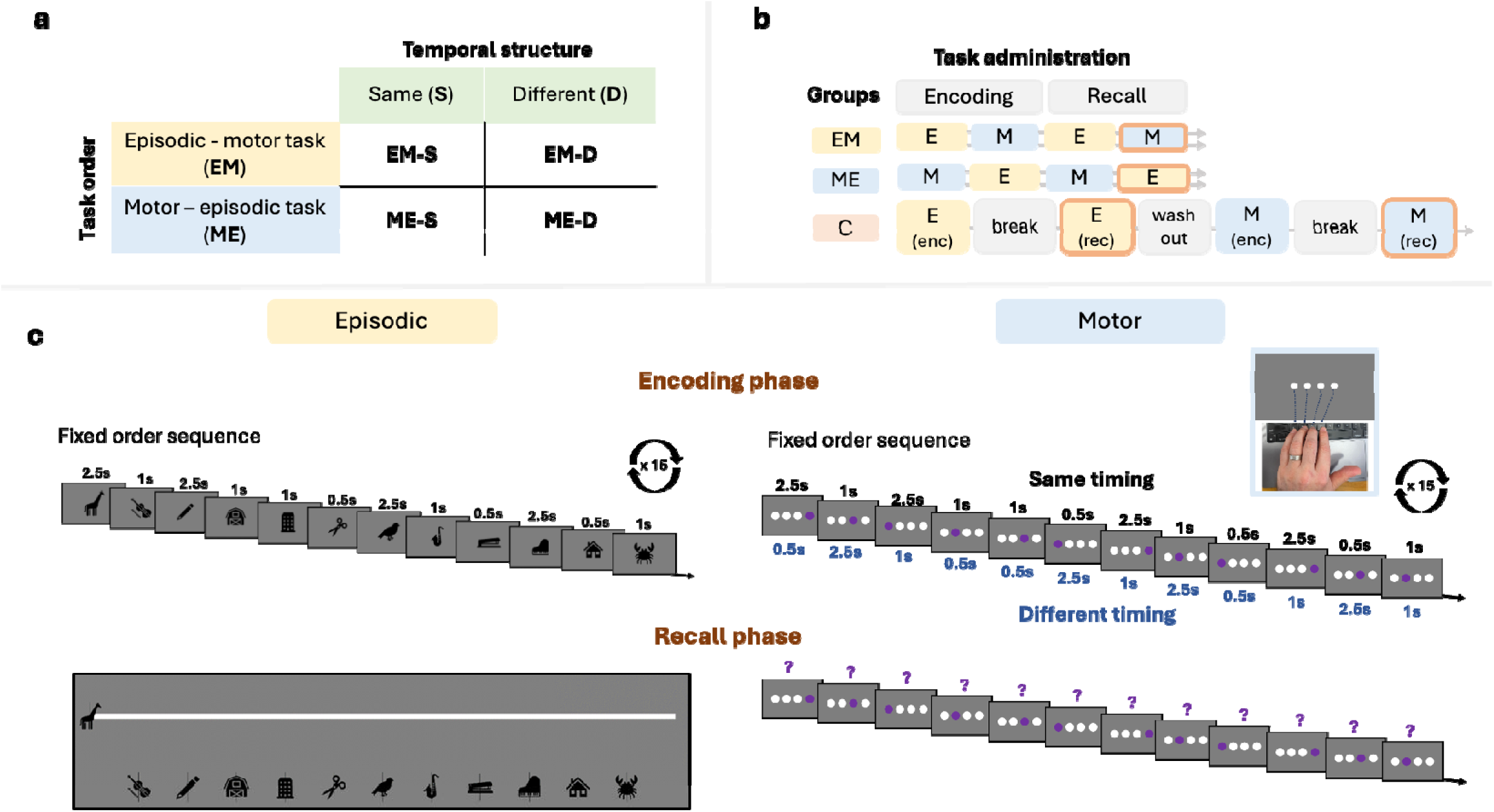
Overview of the study design, protocol and tasks. (a) The study adopted a 2x2 factorial design, with the factors Task order – episodic task followed by the motor task (EM), or the motor task followed by the episodic task (ME) and Temporal structure – same (S) or different (D) timing structures between memory sequences. (b) The order of task administration. Recall phases outlined in orange represent when performance was compared between EM-S and EM-D groups, and between the ME-S and ME-D groups. (c) The encoding (upper panel) and recall phases (lower panel) of the episodic and motor tasks. The temporal structure of the fixed order of images and keypresses during the encoding phase are depicted. For the motor task, the mapping of stimuli to keypress responses is also shown.

Participants in these groups completed the encoding phase of the timing tasks in succession, followed by the recall phase of both tasks (Fig 1b). The temporal tasks reported in this manuscript were completed after a 15-minute washout period during which participants viewed a standardised documentary clip (video: ‘Exploring the Adorable World of Baby Animals’, BBC Earth Kids). Following completion of all tasks, participants were asked if they had actively attended to the timing of stimuli in either task.

In addition, we included a control group (C) (*N* = 20, 4 male) in which participants completed the encoding and recall phases of the episodic and motor task, without an intervening encoding task phase (Fig 1b). For this control group, the encoding and recall phases were separated by a five-to-seven-minute break during which participants engaged in a task-irrelevant activity (origami-making as instructed by a video). The episodic task was administered first to prevent participants from gaining awareness of the need to attend to the timing of the episodic sequence, which may have been difficult to mask had participants completed the motor task first and been probed on the motor keypress timing. The control group was included to capture ‘baseline’ performance on each task (i.e., performance unaffected by the influence of prior or intervening learning).

### Timing tasks

Two novel tasks were developed in PsychoPy (v2023a) (Peirce et al., 2019) to assess participants’ ability to encode and recall the timing of an episodic sequence and a motor sequence. Each task consisted of an encoding phase during which participants viewed or practiced 18 repeats of a timing structure (structures are detailed below), and a recall phase where they reproduced the timing structure. A high number of repetitions (x18) during encoding was adopted to ensure participants developed a schema of the temporal structure (Ghosh & Gilboa, 2014). Prior to recall of the subsequently learned sequence, participants were probed for recall of the first sequence to reactivate this earlier memory to further allow its integration with the subsequently encoded sequence, which has been proposed to give rise to generalisation (Schlichting & Frankland, 2017).

### Episodic task

During the encoding phase, participants viewed a 12-image presentation on a computer screen. Participants were instructed to pay close attention to the presentation as they would be subsequently asked about it in the protocol. To ensure retrospective timing was assessed, participants were not instructed to attend to the timing of the stimuli (MacDonald, 2014; Tsao et al., 2022).

Images were displayed one at a time and were drawn from four categories (animals, stationery, buildings and musical instruments), with three images per category. Across all repeats, the presentation followed a repeating temporal structure (s) with the following durations: 2.5-1-2.5-1-1-0.5-2.5-1-0.5-2.5-0.5-1s (total duration: 16.5s). To mask the temporal nature of the task, the order of images in the first three rounds of the presentation were pseudorandomised (i.e., no images from the same category were presented successively), followed by two foil recognition questions where participants were asked if they saw images that had not been previously displayed in the presentation. In the remaining 15 rounds, the images followed a fixed order and timing (Fig 1c, left upper panel) and concluded with a final foil recognition question of an image not previously shown in the presentation. In addition to masking the temporal nature of the task, the recognition foil questions served to verify that participants paid attention to the image presentation.

During the recall phase, a timeline of the image presentation was displayed to participants. The first image of the fixed order presentation was placed at the start of the timeline (i.e., timepoint zero), with the length of the line representing the total 16.5 seconds duration. The other images were placed below the timeline in the correct order (i.e. as they appeared during the fixed order rounds, Fig 1b, lower panel). Participants were instructed to drag and drop the images on the timeline according to their respective timepoints in the presentation. Where clarification was required, the researcher demonstrated placing images on the timeline to indicate that the duration of the preceding image was represented by the space between image pairs. Participants’ accuracy in reproducing the temporal structure of the image presentation, i.e., the changes in durations between the 11 image pairs of the sequence, was the main metric of this task.

### Motor timing task

During the encoding phase, participants viewed four circles arranged horizontally on a computer screen which corresponded to the keys, ‘Z’, ‘X’, ‘C’ and ‘V’ on a QWERTY keyboard. Participants completed this task with their left (non-dominant) hand (Fig 1c). All circles initially appeared white. Participants were instructed to press the corresponding key when a circle changed colour to purple, and to hold down each keypress until the next circle changed colour. As for the episodic task, participants were not given any instruction to attend to the timing of the stimuli.

Participants practiced a 12-item keypress sequence which either followed the same timing structure as the episodic task (durations of: 2.5-1-2.5-1-1-0.5-2.5-1-0.5-2.5-0.5-1s; 16.5s in total, EM-S and ME-S groups) or a different timing structure (0.5-2.5-1-0.5-0.5-2.5-1-2.5-0.5-1-2.5-1s; 16s in total, EM-D, ME-D, and control groups) (Fig 1c, upper panel).

These structures were aligned in terms of the most salient durations between image pairs to ensure memorability was comparable between structures, such that each contained one repetition and three transitions between the shortest (0.5s) and longest (2.5s) durations. As per the episodic task, the order of keypresses during the first three rounds of this task were pseudorandomised to mask the temporal nature of the task, after which the remaining 15 rounds followed a fixed keypress order and timing (Fig 1c, right panel).

During the recall phase, participants viewed the same four circles on the screen but were now required to reproduce the timing structure of one repeat of the keypress sequence from memory. The visual cues signalling the order of button presses followed the fixed order sequence of the encoding phase. Participants were instructed to hold each key for the same duration as they held the key depressed during encoding. Releasing the key caused the visual display to progress to the next cue in the sequence. Participants were instructed to immediately transition to pressing the next key after the preceding key was released. The main metrics of this task were participants’ accuracy in reproducing the keypress durations in relative (durations as a proportion of the total) and absolute (time in seconds) units.

### Data pre-processing and statistical analysis

Data was pre-processed using custom MATLAB scripts (version R2025a) and analysed statistically in JASP (version 0.95.1) (JASP, 2023).

### Episodic task encoding phase

To verify participants’ attentiveness during the encoding phase of the episodic timing task, we determined their accuracy on the three foil recognition questions presented during encoding.

### Episodic timeline accuracy

To derive timing pattern accuracy, the quotient of image pair durations was calculated for the presented timing structure, i.e., the change in durations between successive image pairs was expressed as a ratio (Image n+1 / Image n, 11 values in total). For example, if Image 1 was presented for 2.5 s and Image 2 was presented for 1 s, then the quotient was 0.4 (Image 2 was 0.4 times the duration of Image 1). Quotients were also derived for the image pair durations produced by the participant at recall. For example, if recall duration of Image 1 was 2.8 s and recall duration of Image 2 was 1.2 s, then the quotient was 0.43 (recalled duration of Image 2 was 0.43 times the duration of Image 1). Accuracy was operationalised as the average of the absolute differences between presented and recalled quotient values.

### Motor timing task encoding phase

We first assessed whether participants modulated the duration of their keypresses during the motor encoding phase according to the presented cue durations. Keypress durations produced by each participant were averaged for each presented cue duration category (0.5s, 1s, 2.5s). Grubbs’ test (□ = .05) was used to assess outliers in the data to detect participants who consistently over-produced keypress durations, relative to other participants in the group.

### Motor timing accuracy

For the recall phase of the motor task, accuracy was quantified by deriving deviation scores that accounted for all possible ordinal representations that participants may have formed during task practice. From the participants’ perspective, there was no clear demarcation between the pseudorandom and fixed order keypresses during encoding. Thus, we aimed to derive metrics that did not penalise participants if they formed a reasonably accurate representation of the timing structure but with an ordinal structure that differed from the fixed order.

For each of the timing structures presented in the motor task, the deviation scores were derived as follows. First, individual keypress durations were calculated as a percentage of the total sequence duration ((duration of keypress *n* / total sequence duration) * 100).

Percentage of the total sequence duration was used instead of absolute keypress duration as inspection of the data revealed that total sequence duration was either compressed or lengthened across participants. These variations in the experience and recall of time have been previously documented in the literature (e.g., Evensmoen et al. (2025); Polti et al. (2022); Press et al. (2014)). Second, quotient values were calculated akin to the episodic timing task metric, i.e., the change in duration between successive keypress durations was expressed as a ratio (Keypress *n*+1 / Keypress *n*, 11 values in total). This resulted in a 12-item percentage sequence, and an 11-item quotient sequence, both as presented to the participant and as produced by the participant.

Next, iterations of these structures were created by shifting the value of the first keypress to all other positions of the sequence (keypresses 2 to 12) by lagging the sequence (i.e. shifting the entire sequence by each ordinal lag, see Figure 2a). The average absolute difference between each lag and the original structure was then obtained. These average values were plotted to form a ‘reference’ lag function based on the presented temporal structure (Figure 2b). These processing steps were also applied to participants’ responses, i.e., average absolute differences between all possible iterations of the structures and participants’ produced structures were obtained to derive a participant function. Finally, deviation scores were obtained by finding the absolute difference between the reference and participant functions. To ensure an unbiased deviation score measure for each participant, the participant function was shifted to align with the minimum point of the reference function prior to determining the average absolute difference (Figure 2c).

**Figure 2.**
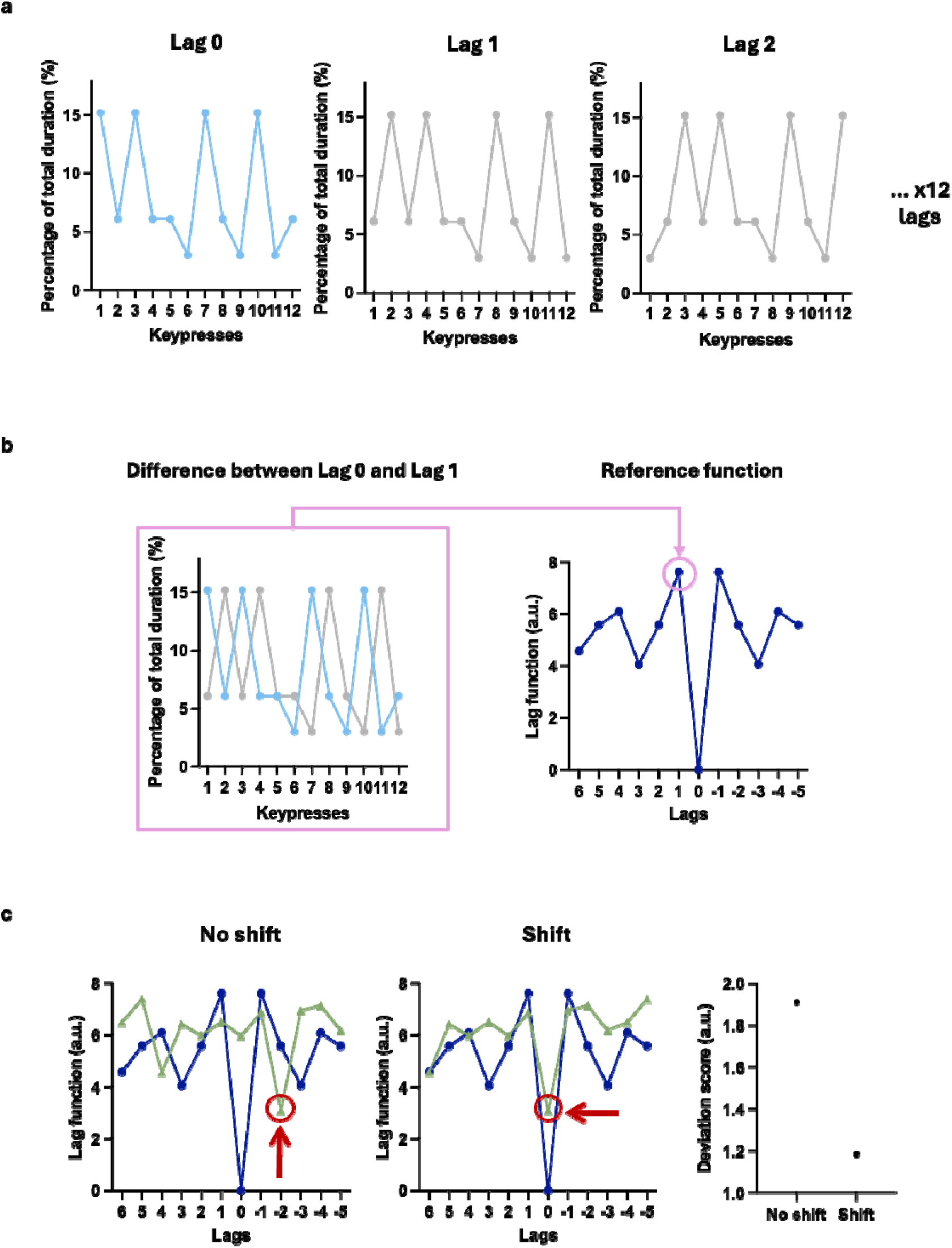
Derivation of deviation scores to quantify motor timing accuracy. These steps were applied to both metrics of motor timing structure (percental of total duration, and quotient values). Percentage of total duration is shown here for simplicity. (a) Iterations of the sequence structure were created by shifting the duration of the first keypress to all other keypress positions in the sequence. To illustrate, Lags 1 and 2 depict sequences where keypress 1 was moved to positions 2 and 3 respectively (forward lag). The original position (Lag 0) is also shown. (b) The average absolute difference between the original (Lag 0) structure and each lag was obtained. These average values were plotted to form the ‘reference’ function. (c) The minimum point of the reference and participant lag functions were aligned before finding the average absolute difference i.e. the deviation score. This shift serves to prevent the deviation score from being artificially inflated, as shown in the rightmost panel. Lower deviation scores reflect better memory of the temporal structure.

We note that we did not analyse participants’ reaction times during this task as initially planned and included in our pre-registration. This choice was decided upon after data collection as reaction times did not appear appropriate to measure motor timing in the context of this task.

### Statistical analyses

We conducted analyses of variance (ANOVA) with the factors of Group (same timing, different timing, and control) for measures of episodic and motor timing accuracy. We applied a Bonferroni correction to between-groups post-hoc comparisons (i.e., α = 0.05/3 = 0.017). Bayes factors were also determined (Bayesian ANOVA and independent *t*-test, Cauchy parameter = 0.707) to establish the relative evidence in favour of the null or alternative hypothesis (Ly et al., 2016; Rouder et al., 2009). BF_10_ values are reported for significant results to indicate the strength of evidence for the alternative hypothesis (i.e., that generalisation has occurred), while BF_01_ values are reported for non-significant results to indicate strength of evidence for the null hypothesis. For both, values ranging from .33 to 3 is considered weak evidence, 3 to 10 moderate evidence and values larger than 10 being strong evidence (van Doorn et al., 2021).

To explore relationships between motor and episodic timing recall, Pearsons’ correlations were computed between the accuracy metrics of the motor and episodic tasks.

### Data exclusions

Participant data were excluded from recall phase analyses if the participant did not complete the encoding and recall phases of a given task correctly (please see Supplemental File for detailed exclusion criteria by task phase). Additionally, as we were interested in examining cross-domain transfer effects, we also excluded participants from analyses if they did not complete the first task correctly. To illustrate how these criteria were applied, in the case of motor-to-episodic transfer, a participant would be excluded if they did not complete the encode the motor timing structure as intended, and/or if they met the exclusion criteria for the episodic task. A breakdown of participant exclusions by group and associated reasons are reported in the Supplemental File (range, 2 to 5 participants per group).

## Results

### Participants attended to the image presentation during the episodic task encoding phase

Across the sample, one participant (EM-S group) responded ‘yes’ to all episodic image foil questions (as per exclusion criteria described in the Supplemental File, this participant was removed from subsequent motor analyses). Overall, 91.61% of participants (across all groups) responded accurately to all foil questions, indicating that participants attended to the image presentation.

### Participants modulated keypress durations according to cue durations during the motor task encoding phase

Across all groups, participants demonstrated a tendency to produce keypresses that were approximately 400ms longer than the durations of the coloured circle cues during the motor encoding phase (Figure 3). Specifically, the average produced duration keypresses across groups were .97s, 1.43, and 2.91s in length (vs. .5s, 1s, 2.5s cue durations).

**Figure 3.**
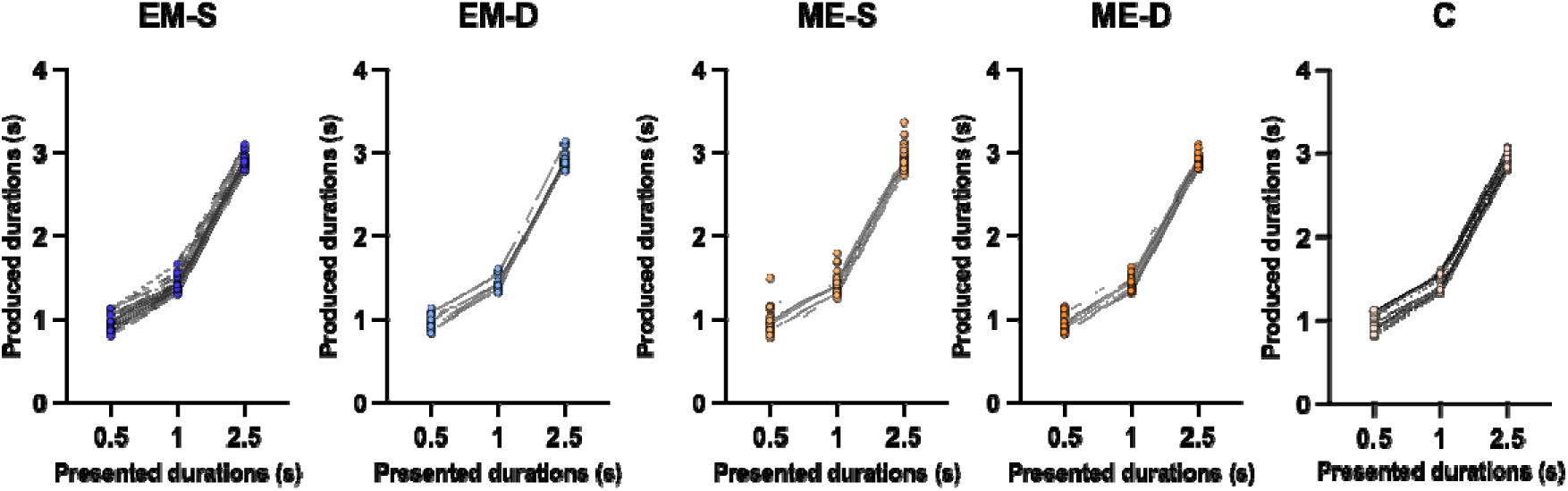
Keypress durations produced by participants during the encoding phase of the motor task, per group. Each line represents an individual participant. An outlier was detected in the ME-S group, over-producing the keypress durations as depicted in the graph.

Follow-up analyses revealed that the timing percentages and quotient values produced by participants during encoding were qualitatively similar to the cued durations, suggesting that the intended manipulation of timing structures was achieved (Figure 4).

**Figure 4.**
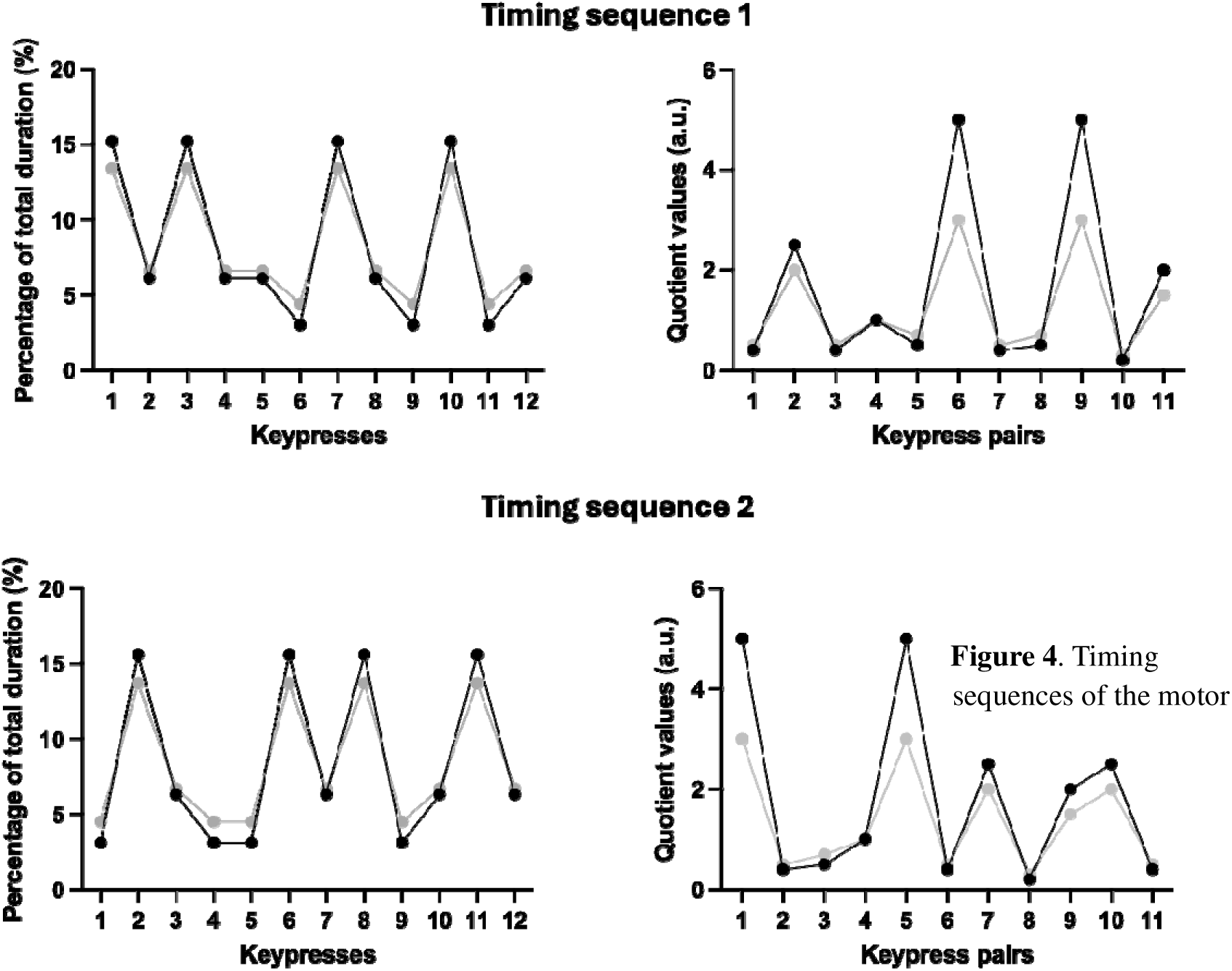
Timing sequences of the motor encoding phase. The black lines represent the keypress durations that were cued to participants. The grey lines represent structures constructed from the average keypress durations participants produced during encoding (i.e., .97s, 1.43, and 2.91s). Timing sequence 1 was practiced by participants in the EM-S and ME-S groups (same timing across tasks), while Timing sequence 2 was practiced by participants in the EM-D, ME-D and C groups (different timings across tasks).

### Recall of the first memory did not differ across groups

When the motor sequence was encoded first, ANOVA revealed no differences in motor timing accuracy between ME-S, ME-D and control groups (proportion metrics: *p =* .425, BF_01_ = 4.01; pattern metrics: *p* = .474, BF_01_ = 3.99). A similar pattern was observed for episodic recall accuracy, which did not differ between the EM-S, EM-D and control groups (*p* = .806, BF_01_ = 7.27). These findings suggest that the representation of the first memory was not affected by the intervening encoding phase and was thus likely recalled / reactivated with fidelity.

### No evidence of generalisation of timing information from the episodic to motor domain

We hypothesised that a shared timing structure between a motor and episodic sequence would facilitate reproduction of the motor timing sequence. Contrary to expectations, we observed no main effect of Group on participants’ accuracy in reproducing the timing of the motor sequence. This null effect was present regardless of the metric used to examine accuracy – either in terms of keypress durations as a percentage of total duration (*M*_EM-S_ = 1.36, *SD*_EM-S_ = .38; *M*_EM-D_ = 1.33, *SD*_EM-D_ = .24; M_C_ = 1.33, SD_C_ = .3) (*F* (2, 67) = .063, *p* = .939, η^2^_p_ = .002, BF_01_ = 7.75), or for the quotient value timing pattern (*M*_EM-S_ = .52, *SD*_EM-S_ = .099; *M*_EM-D_ = .52, *SD*_EM-D_ = .10; M_C_ = .49, SD_C_ = .096) (*F* (2, 66) = .53, *p* = .59, η^2^_p_ = .016, BF_01_ = 5.47) (Figure 5). This implies that motor timing accuracy did not differ between groups who had first encoded an episodic sequence with the same versus different timing.

**Figure 5.**
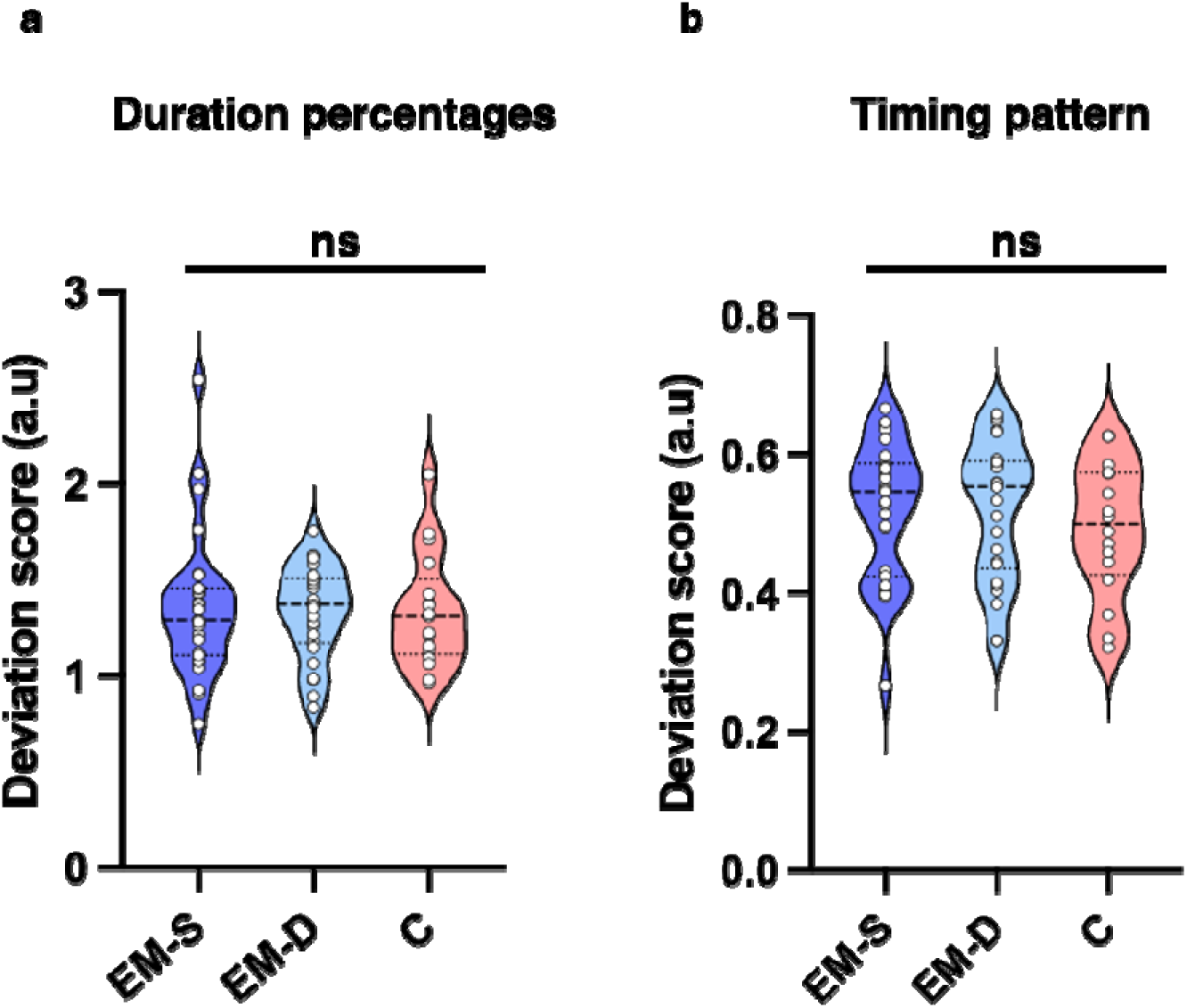
Participants accuracy in reproducing the timing of the motor sequence. No significant differences were documented in accuracy rates between groups.

### Prior motor learning interfered with recall of episodic timing

There was a main effect of Group on episodic timing accuracy at recall (*F* (2, 70) = 3.492, *p* = .036, η^2^_p_ = .091). Unexpectedly, post-hoc tests revealed that accuracy was poorer for participants who first learned a motor sequence with the same (*M*_ME-S_ = 1.35, *SD*_ME-S_ = .15) compared to different (*M*_ME-D_ = 1.25, *SD*_ME-D_ = .16) timing as the episodic sequence, *t* (52) = -2.53, *p* = .014, *d* = .68, 95%CI*_d_* = [1.23,.14], BF_10_ = 3.603 (Fig 6a). Accuracy did not differ between the control group (M_C_ = 1.25, SD_C_ = .18) and participants who first learned a motor sequence with the same timing (*t* (42) = -2.04, *p* = .047, *d* = .63, 95%CI*_d_* = [1.24, .007], BF_01_ = .648), or different timing (*t* (45) = .08, *p* = .939, *d* = .023, 95%CI*_d_* = [-.565, .611], BF_01_ = 3.366). These findings suggest that prior learning of a motor sequence may interfere with subsequent recall of an episodic timeline when both sequences share a timing structure. The timing pattern presented to participants alongside the average timing patterns produced by each group are depicted in Fig 6b for visualisation purposes.

**Figure 6.**
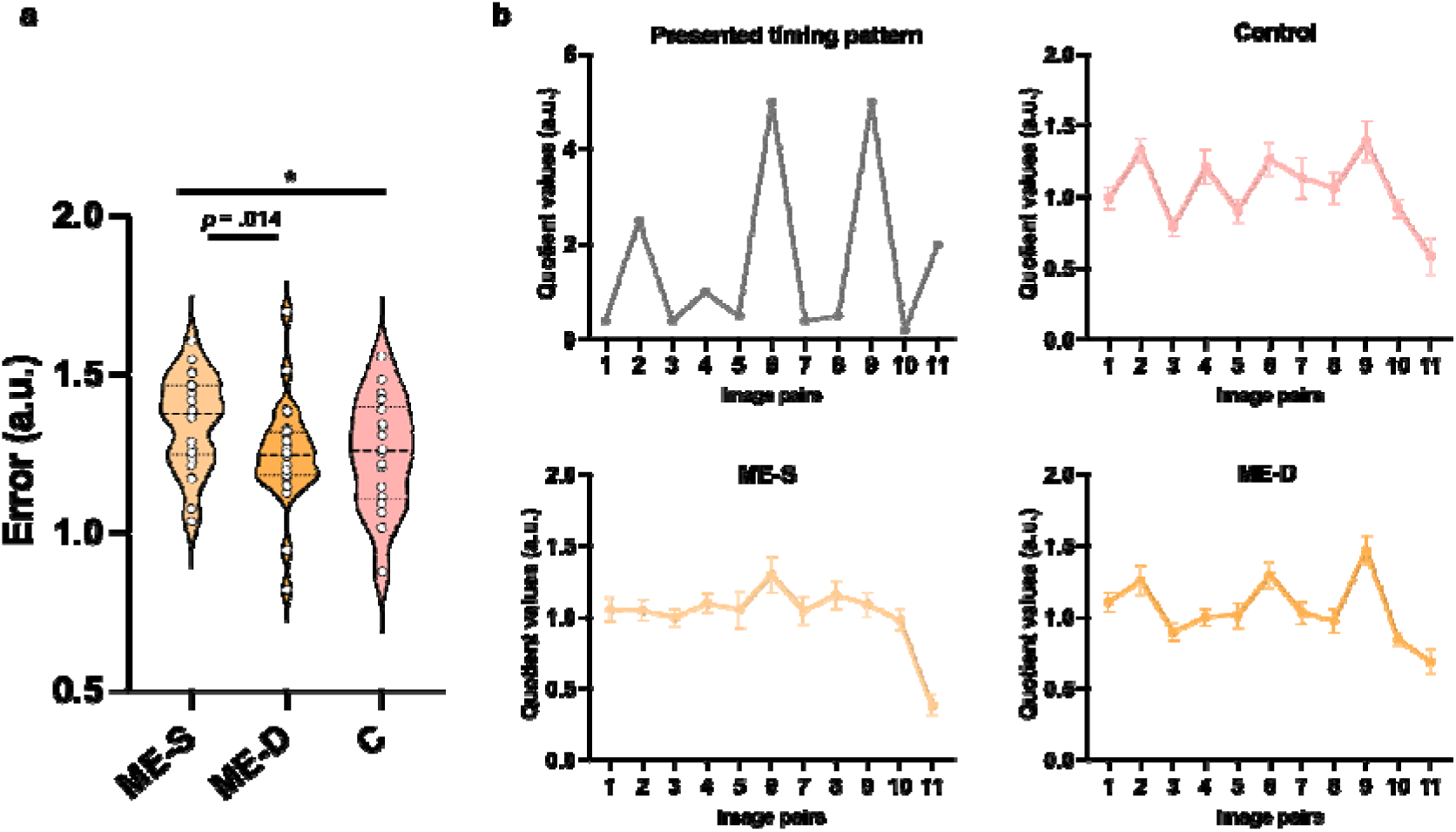
(a) Accuracy in reproducing the timeline of the episodic sequence across all groups. Participants in the same timing group demonstrated poorer recall than participants in the different timing group. (b) Visualisation of the episodic timelines produced by each group. Each point represents the quotient value averaged across participants within a group. Error bars represent ± SE. * *p* < .05.

### Correlations between episodic and motor timing recall

No correlations were observed between motor and episodic accuracy metrics across all groups (all *p*s > .1).

## Discussion

The present study examined generalisation between motor and episodic memory sequences with a shared timing structure. Contrary to expectations, reproduction of a motor timing sequence was not influenced by the temporal structure of an episodic sequence learned beforehand. Furthermore, recall of an episodic sequence timeline was significantly worse for participants who had previously learned a motor sequence with the same timing structure.

This finding suggests that prior learning resulted in interference as opposed to generalisation (facilitation) of the subsequent memory. Despite the absence of generalisation effects, the finding of cross-memory interference suggests an interaction and/or overlap in the neural underpinnings that encode time across our different memory systems.

The interference of episodic recall by a previously learned motor sequence with a shared timing structure may reflect involvement of overlapping neural ensembles encoding the structure of distinct memories (Herszage & Censor, 2018; Robertson, 2018). From a neurophysiological perspective, interference as opposed to generalisation may reflect state-dependency of the neural ensembles encoding temporal structure across memories (Robertson, 2018). Cross-memory generalisation is theorised to occur when the neuronal ensembles involved in encoding the first memory, whilst still excitable, become allocated to processing the same structure encountered in a subsequent memory / experience (Robertson, 2022). An increase in excitability accompanies the formation of new memories, while a gradual decrease allows memories to stabilise (Barron, 2021; Barron et al., 2017).

Stabilisation generally occurs on the timescale of hours (Mosha & Robertson, 2016), however, repeated practice of a visual discrimination task (to the point of overlearning) has been shown to rapidly hyper-stabilise the developed skill (Shibata et al., 2017). While participants were not trained to overlearn the motor sequence in this study, repeated practice of the motor sequence may have stabilised, to an extent, the ensembles that encoded the preceding temporal structure. Indeed, concurrent actions during memory encoding have been shown to strengthen memory traces (Yebra et al., 2019). Hypothetically, the stability of the formed representation may have reduced the capacity for subsequent memory allocation and, by extension, the ability for generalisation to occur.

An alternate explanation can be drawn from the literature on anterograde interference within the motor learning domain. Anterograde interference occurs when prior learning interferes with subsequent learning, which suggests that the capacity of a memory system to undergo plastic changes is limited after initial learning (Cantarero et al., 2013; Hamel et al., 2022). Anterograde interference is graded by the similarity between memories and the magnitude of prior learning, where greater similarity and prior practice evoke greater interference (De La Fontaine et al., 2023; Hamel et al., 2022), while greater interference has also been correlated with greater occlusion of plasticity following initial learning (Cantarero et al., 2013). As the present interference effect was documented on the basis of a shared structure, it is possible that plasticity across regions encoding for the shared temporal structure was temporarily occluded following initial learning.

In the broader framework of memory systems, the finding of interference contributes to the growing body of work showing that motor and declarative systems may share greater similarities than traditionally appreciated, which to-date is comprised largely of findings on the ordinal aspect of memories (e.g., Jacobacci et al. (2020); Kim and Lee (2023); Yewbrey and Kornysheva (2024)). Specifically, this finding suggests that timing information such as the regularities or patterns of durations in each sequence (Sherman et al., 2020) may be encoded by similar neural regions across systems. Learning probabilistic regularities of sequential stimuli is generally deemed to be hippocampal-dependent (Schapiro & Turk-Browne, 2015; Sherman et al., 2020), while temporal regularities have been shown to be represented within the supplementary motor area (Otsuka & Saiki, 2025). Interestingly, this finding appears to align with theories positing that the motor system encodes timing information across domains, and may interact with other brain regions relevant for a given task context (Merchant et al., 2013; Sanchez-Moncada et al., 2024). Future work using non-invasive brain stimulation to modulate regions such as the supplementary motor area or hippocampus may help delineate the locus of cross-domain memory interference.

We did not observe any influence of prior episodic encoding on subsequent motor timing recall, nor correlations between task metrics. The reasons for this are not immediately clear, however, in relation to the former finding, based on theories of neuronal excitability discussed above, it is possible that the representation of the motor sequence participants developed was stable and thus robust to prior and subsequent learning. Alternatively, the measures derived to quantify motor accuracy may not have been adequately sensitive. This factor may have also contributed to the lack of correlations between motor and episodic metrics. However, the motor recall metrics were derived with the goal of accounting for any misattributions of keypress durations and thus represent unbiased measures of motor timing accuracy.

A potential limitation of this study lies in the protocol of task administration; specifically, the inclusion of the recall phase of the first memory prior to the recall phase of the second memory. This initial recall phase was included with the rationale of reactivating the memory of the first sequence as a cue that may potentially facilitate recall of the subsequently learned sequence when both sequences shared a structure (Schlichting & Frankland, 2017). Indeed, reactivated memories can influence subsequently learned information (Tambini & Davachi, 2019). However, any potential positive effects of reactivation are predicated on the assumption that participants’ memory for the first sequence was fairly accurate, while reactivated memories are at the risk of becoming disrupted upon reactivation (Lee et al., 2017). Nonetheless, the current protocol provides an ecologically valid paradigm reflecting how several competing memories are generally encoded and recalled in daily life. Future work examining memory interactions may perhaps assess recall of the subsequent memory without a preceding recall of the first memory.

In conclusion, we showed that a shared timing structure with a preceding motor sequence led to poorer recall of an episodic timeline - i.e., interference as opposed to generalisation or facilitated recall of the subsequent memory as originally hypothesised. Nonetheless, the finding of interference provides behavioural evidence that aligns with neuroimaging studies suggesting that that timing information may be encoded by shared neuronal ensembles across motor and declarative systems.

## General Disclosures

### Artificial intelligence

No artificial intelligence was used in the writing or production of this manuscript.

### Ethics

Ethical approval was obtained from the Monash University Human Research Ethics Committee

## References

Barron, H. C. (2021). Neural inhibition for continual learning and memory. Curr Opin Neurobiol, 67, 85–94. 10.1016/j.conb.2020.09.007

Barron, H. C., Vogels, T. P., Behrens, T. E., & Ramaswami, M. (2017). Inhibitory engrams in perception and memory. Proc Natl Acad Sci U S A, 114(26), 6666–6674. 10.1073/pnas.1701812114

Breska, A., & Ivry, R. B. (2016). Taxonomies of Timing: Where Does the Cerebellum Fit In? Curr Opin Behav Sci, 8, 282–288. 10.1016/j.cobeha.2016.02.034

Cantarero, G., Tang, B., O’Malley, R., Salas, R., & Celnik, P. (2013). Motor learning interference is proportional to occlusion of LTP-like plasticity. J Neurosci, 33(11), 4634–4641. 10.1523/JNEUROSCI.4706-12.2013

De La Fontaine, E., Hamel, R., Lepage, J. F., & Bernier, P. M. (2023). The influence of learning history on anterograde interference. Neurobiol Learn Mem, 206, 107866. 10.1016/j.nlm.2023.107866

Evensmoen, H. R., Rimol, L. M., Rise, H. S., Hansen, T. I., Nili, H., Winkler, A. M., & Haberg, A. (2025). Pattern integration and differentiation: Dual process model of episodic memory. Imaging Neurosci (Camb), 3. 10.1162/imag_a_00433

Failla, A., Bracco, M., & Robertson, E. M. (2025). A common serial structure causes offline excitability changes linked to generalization between different memory types. Curr Biol. 10.1016/j.cub.2025.10.063

Ghosh, V. E., & Gilboa, A. (2014). What is a memory schema? A historical perspective on current neuroscience literature. Neuropsychologia, 53, 104–114. 10.1016/j.neuropsychologia.2013.11.010

Guo, J., Zhang, Z., Makwana, M., Sternad, D., & Song, J. H. (2025). Emergent motor timing enhances time perception. iScience, 28(9), 113367. 10.1016/j.isci.2025.113367

Hamel, R., Lepage, J. F., & Bernier, P. M. (2022). Anterograde interference emerges along a gradient as a function of task similarity: A behavioural study. Eur J Neurosci, 55(1), 49–66. 10.1111/ejn.15561

Henke, K. (2010). A model for memory systems based on processing modes rather than consciousness. Nat Rev Neurosci, 11(7), 523–532. 10.1038/nrn2850

Herszage, J., & Censor, N. (2018). Modulation of Learning and Memory: A Shared Framework for Interference and Generalization. Neuroscience, 392, 270–280. 10.1016/j.neuroscience.2018.08.006

Jacobacci, F., Armony, J. L., Yeffal, A., Lerner, G., Amaro, E., Jr., Jovicich, J., Doyon, J., & Della-Maggiore, V. (2020). Rapid hippocampal plasticity supports motor sequence learning. Proc Natl Acad Sci U S A, 117(38), 23898–23903. 10.1073/pnas.2009576117

JASP. (2023). JASP (Version 0.17.2.1) [Computer Software]. In

Kim, J. S., & Lee, S. A. (2023). Hippocampal orchestration of associative and sequential memory networks for episodic retrieval. Cell Rep, 42(8), 112989. 10.1016/j.celrep.2023.112989

Kornysheva, K., Sierk, A., & Diedrichsen, J. (2013). Interaction of temporal and ordinal representations in movement sequences. J Neurophysiol, 109(5), 1416–1424. 10.1152/jn.00509.2012

Lee, J. L. C., Nader, K., & Schiller, D. (2017). An Update on Memory Reconsolidation Updating. Trends Cogn Sci, 21(7), 531–545. 10.1016/j.tics.2017.04.006

Ly, A., Verhagen, J., & Wagenmakers, E. J. (2016). Harold Jeffreys’s default Bayes factor hypothesis tests: Explanation, extension, and application in psychology. Journal of Mathematical Psychology 72, 19–32. 10.1016/j.jmp.2015.06.004

MacDonald, C. J. (2014). Prospective and retrospective duration memory in the hippocampus: is time in the foreground or background? Philos Trans R Soc Lond B Biol Sci, 369(1637), 20120463. 10.1098/rstb.2012.0463

Meegan, D. V., Aslin, R. N., & Jacobs, R. A. (2000). Motor timing learned without motor training. Nat Neurosci, 3(9), 860–862. 10.1038/78757

Merchant, H., Harrington, D. L., & Meck, W. H. (2013). Neural basis of the perception and estimation of time. Annu Rev Neurosci, 36, 313–336. 10.1146/annurev-neuro-062012-170349

Mosha, N., & Robertson, E. M. (2016). Unstable Memories Create a High-Level Representation that Enables Learning Transfer. Curr Biol, 26(1), 100–105. 10.1016/j.cub.2015.11.035

Mutanen, T. P., Bracco, M., & Robertson, E. M. (2020). A Common Task Structure Links Together the Fate of Different Types of Memories. Curr Biol, 30(11), 2139–2145 e2135. 10.1016/j.cub.2020.03.043

Otsuka, S., & Saiki, J. (2025). Neural representations of visual statistical learning based on temporal duration. Imaging Neurosci (Camb), 3. 10.1162/IMAG.a.135

Peirce, J., Gray, J. R., Simpson, S., MacAskill, M., Hochenberger, R., Sogo, H., Kastman, E., & Lindelov, J. K. (2019). PsychoPy2: Experiments in behavior made easy. Behav Res Methods, 51(1), 195–203. 10.3758/s13428-018-01193-y

Polti, I., Nau, M., Kaplan, R., van Wassenhove, V., & Doeller, C. F. (2022). Rapid encoding of task regularities in the human hippocampus guides sensorimotor timing. Elife, 11. 10.7554/eLife.79027

Press, C., Berlot, E., Bird, G., Ivry, R., & Cook, R. (2014). Moving time: the influence of action on duration perception. J Exp Psychol Gen, 143(5), 1787–1793. 10.1037/a0037650

Robertson, E. M. (2018). Memory instability as a gateway to generalization. PLoS Biol, 16(3), e2004633. 10.1371/journal.pbio.2004633

Robertson, E. M. (2022). Memory leaks: information shared across memory systems. Trends Cogn Sci, 26(7), 544–554. 10.1016/j.tics.2022.03.010

Rouder, J. N., Speckman, P. L., Sun, D., Morey, R. D., & Iverson, G. (2009). Bayesian t tests for accepting and rejecting the null hypothesis. Psychon Bull Rev, 16(2), 225–237. 10.3758/PBR.16.2.225

Sanchez-Moncada, I., Concha, L., & Merchant, H. (2024). Pre-supplementary Motor Cortex Mediates Learning Transfer from Perceptual to Motor Timing. J Neurosci, 44(8). 10.1523/JNEUROSCI.3191-20.2023

Schapiro, A. C., & Turk-Browne, N. B. (2015). Statistical Learning. In A. Toga (Ed.), Brain Mapping: An Encyclopedic Reference (Vol. 3, pp. 501–506): Academic Press: Elsevier.

Schlichting, M. L., & Frankland, P. W. (2017). Memory allocation and integration in rodents and humans. Current Opinion in Behavioral Sciences, 17, 90–98. 10.1016/j.cobeha.2017.07.013

Sherman, B. E., Graves, K. N., & Turk-Browne, N. B. (2020). The prevalence and importance of statistical learning in human cognition and behavior. Curr Opin Behav Sci, 32, 15–20. 10.1016/j.cobeha.2020.01.015

Shibata, K., Sasaki, Y., Bang, J. W., Walsh, E. G., Machizawa, M. G., Tamaki, M., Chang, L. H., & Watanabe, T. (2017). Overlearning hyperstabilizes a skill by rapidly making neurochemical processing inhibitory-dominant. Nat Neurosci, 20(3), 470–475. 10.1038/nn.4490

Squire, L. R. (2004). Memory systems of the brain: a brief history and current perspective. Neurobiol Learn Mem, 82(3), 171–177. 10.1016/j.nlm.2004.06.005

Tambini, A., & Davachi, L. (2019). Awake Reactivation of Prior Experiences Consolidates Memories and Biases Cognition. Trends Cogn Sci, 23(10), 876–890. 10.1016/j.tics.2019.07.008

Tsao, A., Yousefzadeh, S. A., Meck, W. H., Moser, M. B., & Moser, E. I. (2022). The neural bases for timing of durations. Nat Rev Neurosci, 23(11), 646–665. 10.1038/s41583-022-00623-3

van Doorn, J., van den Bergh, D., Bohm, U., Dablander, F., Derks, K., Draws, T., Etz, A., Evans, N. J., Gronau, Q. F., Haaf, J. M., Hinne, M., Kucharsky, S., Ly, A., Marsman, M., Matzke, D., Gupta, A., Sarafoglou, A., Stefan, A., Voelkel, J. G., & Wagenmakers, E. J. (2021). The JASP guidelines for conducting and reporting a Bayesian analysis. Psychon Bull Rev, 28(3), 813–826. 10.3758/s13423-020-01798-5

Yebra, M., Galarza-Vallejo, A., Soto-Leon, V., Gonzalez-Rosa, J. J., de Berker, A. O., Bestmann, S., Oliviero, A., Kroes, M. C. W., & Strange, B. A. (2019). Action boosts episodic memory encoding in humans via engagement of a noradrenergic system. Nat Commun, 10(1), 3534. 10.1038/s41467-019-11358-8

Yewbrey, R., & Kornysheva, K. (2024). The Hippocampus Preorders Movements for Skilled Action Sequences. J Neurosci, 44(45). 10.1523/JNEUROSCI.0832-24.2024

