## Supplemental file for "Investigating cross-memory interactions of timing structures"

### Data exclusion criteria by tasks and task phase

*Episodic encoding phase*

1) Answered all recognition foil questions incorrectly, suggesting that they did not attend to the image presentation to a sufficient degree
2) Actively attended to the timing of the images, inferred from participants self-reporting the expectation they would be asked about the timing of the image presentation

*Episodic recall phase*

1) Participants self-reporting that they spaced the images on the timeline to equate the distances between images, vs. according to the duration of the images
2) Outliers in error rates as detected by Grubbs test ($\alpha = .05$)

*Motor encoding phase*

1) Consistent over-production of keypress durations during the encoding phase of the motor task, suggesting an inability to learn the motor timing structure, as detected by Grubbs test ($\alpha = .05$)

*Motor recall phase*

1) Variances in produced keypress durations that were below .1 or 3SD above the average variance for a given group (i.e., very low or very high variance),
2) Reproduced keypresses that were consistently around reaction times (~350ms) and/or
3) Outliers in deviation scores data as detected by Grubbs test ($\alpha = .05$)

For tests of transfer effects, data were additionally excluded where participants did not complete the encoding phase of the preceding task as intended. The total number of exclusions based on these criteria  per group are reported in Supplementary Figure 1.

**Episodic to motor transfer (motor recall phase)**

**EM-S (*N* = 30)**

**Percentage deviation score**
Total *N* excluded = 4
- Outlier (1)
- Keypress variance at recall <.1 (1)
- Attended to timing of preceding episodic sequence (1)
- Misunderstood task instructions (1)

**Pattern deviation score**
Total *N* excluded = 4
- Outlier (1)
- Keypress variance at recall <.1 (1)
- Attended to timing of preceding episodic sequence (1)
- Misunderstood task instructions (1)

**EM-D (*N* = 30)**

**Percentage deviation score**
Total *N* excluded = 4
- Keypress durations at recall within average reaction time range (1)
- Keypress variance at recall > 3SD of average variance (1)
- Attended to timing of preceding episodic sequence (2)

**Pattern deviation score**
Total *N* excluded = 4
- Keypress durations at recall within average reaction time range (1)
- Keypress variance at recall > 3SD of average variance (1)
- Attended to timing of preceding episodic sequence (2)

**Motor to episodic transfer (episodic recall phase)**

**ME-S (*N* = 30)**

**Pattern error score**
Total *N* excluded = 4
- Outlier (1)
- Actively attended to episodic sequence timing (2)
- Did not complete motor encoding phase correctly (1)

**ME-D (*N* = 33)**

**Pattern error score**
Total *N* excluded = 4
- Actively attended to episodic sequence timing (3)
- Did not complete motor encoding phase correctly (1)

**Control (*N* = 20)**

**Pattern error score (episodic recall)**
Total *N* excluded = 2
- Did not place images on timeline according to durations (2)

**Percentage deviation score (motor recall)**
Total *N* excluded = 3
- Outliers (2)
- Keypress variance at recall > 3SD of average variance (1)

**Pattern deviation score (motor recall)**
Total *N* excluded = 4
- Outliers (3)
- Keypress variance at recall > 3SD of average variance (1)

**Recall data of first memory**

**EM-S**
**Pattern error score (episodic recall)**
Total *N* excluded = 1
- Actively attended to episodic sequence timing (1)

**EM-D**
**Pattern error score (episodic recall)**
Total *N* excluded = 2
- Actively attended to episodic sequence timing (2)

**ME-S**
**Percentage and pattern deviation scores (motor recall) [same participant exclusions for both metrics]**
Total *N* excluded = 7
- Outlier (1)
- Keypress durations at recall within average reaction time range (5)
- Keypress variance at recall <.1 (1)

**ME-D**
**Percentage deviation scores (motor recall)**
Total *N* excluded = 6
- Keypress durations at recall within average reaction time range (3)
- Keypress variance at recall <.1 (2)
- Did not complete motor encoding phase correctly (1)

**Pattern deviation scores (motor recall)**
Total *N* excluded = 9
- Outliers (3)
- Keypress durations at recall within average reaction time range (3)
- Keypress variance at recall <.1 (2)
- Did not complete motor encoding phase correctly (1)

**Supplementary Figure 1.** Data exclusions per group for motor and episodic recall phases.
